# Computer Assisted Localization of a Heart Arrhythmia

**DOI:** 10.1101/365932

**Authors:** Chris Vogl, Peng Zheng, Stephen P. Seslar, Aleksandr Y. Aravkin

## Abstract

We consider the problem of locating a point-source heart arrhythmia using data from a standard diagnostic procedure, where a reference catheter is placed in the heart, and arrival times from a second diagnostic catheter are recorded as the diagnostic catheter moves around within the heart.

We model this situation as a nonconvex feasibility problem, where given a set of arrival times, we look for a source location that is consistent with the available data. We develop a new optimization approach and fast algorithm to obtain online proposals for the next location to suggest to the operator as she collects data. We validate the procedure using a Monte Carlo simulation based on patients’ electrophysiological data. The proposed procedure robustly and quickly locates the source of arrhythmias without any prior knowledge of heart anatomy.

## I. Introduction

Catheter ablation is the treatment of choice to diagnose and treat cardiac arrhythmias. Accurately determining the origin of a cardiac arrhythmia is of critical importance in catheter ablation procedures. In many instances, arrhythmias originate from a focal point source and the electrical signal spreads concentrically in all directions away from that point. The traditional method of localizing such arrhythmias involves a user-directed movement of a mapping catheter through a cardiac chamber — a somewhat haphazard “hunt and peck” method. The arrhythmia signal arrival time on the roving map catheter is compared against the signal arrival time on a stationary reference catheter until the *earliest relative timing* site is identified.

Based on inherent limitations in how humans recognize patterns, this process requires a significant area of the heart chamber to be mapped before the operator can start to hone in on the location of the arrhythmia source. The operator is essentially required to solve an optimization problem by hand to minimize the arrival time relative to that obtained by a stationary catheter.

We propose a computer-assisted mapping system that can effectively use all available information and point the operator to the ‘next touch’ location. This approach can substantially reduce the time and number of touch points needed to reliably determine the site of arrhythmia origin. We develop a method for arrhythmia localization and evaluate it using Monte Carlo simulations based on a deidentified electroanatomic map from a patients that underwent mapping and acutely successful ablation of a focal arrhythmia using the Rhythmia mapping system^1^.

**Related work**. We found one prior approach to localizing arrhythmia using optimization [5]. The approach is similar in principle, and uses optimization to recommend the next point to sample by the operator. The key differentiating factor is that [5] assumes the existence of an anatomical map; the approach is predicated on being able to solve a linear regression problem that uses *every available nodal point* as a potential origin, and then pick the one that best fits the available data. In contrast, we make no assumptions about the existence of anatomical maps; our proposals are based only on observations taken by the operator, and we can help patients who have undergone no prior mapping studies. In addition, the algorithm of [5], for each recommendation, must solve a number of regression problems equal to the number of points in their mesh. In contrast, we only need to solve one feasibility problem to find the next point proposal.

The paper proceeds as follows. In Section II, we formulate arrhythmia localization as a feasibility problem and derive an associated optimization problem. In Section III we develop a fast algorithm, with practical considerations for dealing with noisy data and online application of arrhythmia localization. In Section IV we present results of the approach using patient arrival time data, and end with conclusions in Section V.

## II. Formulation

We are given a set of (ordered) relative arrival times

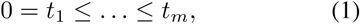

obtained by finding the differences between times recorded using a reference catheter and a diagnostic catheter at 3D locations *x*_1;_…, *x_m_* within the heart.

**Simple Feasible Region**. We look for a source location *x_s_* consistent with available observations. We assume that the local signal transmission speed s around the source *x_s_* is known and given. Denoting the actual arrival times by 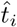, we get relations

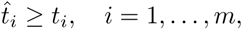

which translate to the constraints

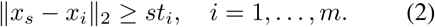

Finding a source that is consistent with the available observations is equivalent to finding *x_s_* that satisfies (2). The feasible region from (2) is shown in panel (a) of Figure 1; the true source must lie outside of the union of the disks shown in the figure.

**Fig. 1.**
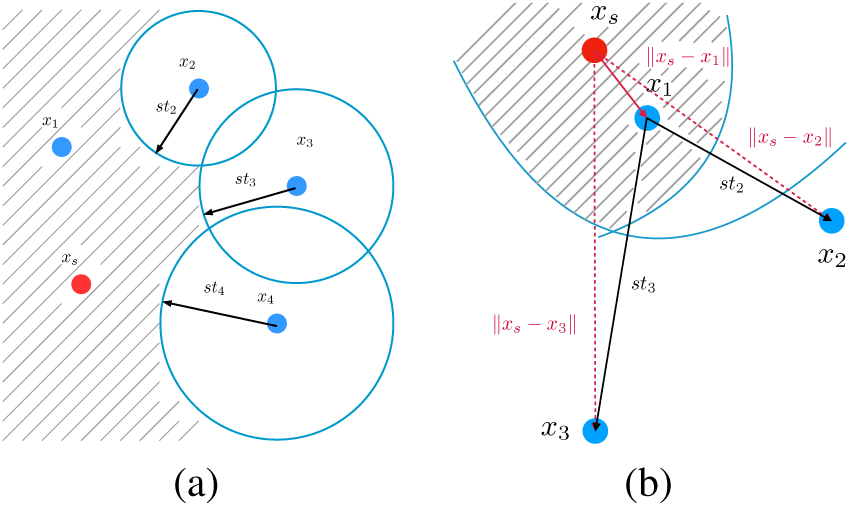
Simple timing relationships define a feasibility problem for locating the source location *x_s_* of the arrhythmia. (a) The set Ω_1_ described by (2). (b) The set Ω_2_ described by (3).

**Coupled Feasible Region**. A more powerful formulation incorporates first arrival information, similar to the fast marching method used to solve the Eikonal equation [4]. We look for *x_s_* satisfying

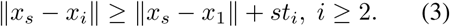

The fast marching method and the inequalities (3) enforce a reverse triangle inequality: the time the signal takes to go from *x_s_* to *x_j_* is is larger than the time needed to go from *x_s_* to *x*_1_ and then *x*_1_ to *x_i_*; otherwise we would not have observed the given distribution of arrival times. The resulting region is shown in panel (b) of Figure 1.

## III. Algorithm

Nonconvex feasibility problems can be solved using optimization techniques such as alternating optimization methods or Douglas-Rachford splitting [2]. These algorithms can take hundreds to thousands of iterations for simple problem instances [2, Table 1]. We propose a new relaxation that converges very rapidly, generating a feasible solution *x_s_* within a few iterations in most instances.

In developing relaxations for (2) and (3), we use the ideas recently developed by [6]. We introduce auxiliary variables *w_i_* to approximate each *x_s_* − *x_i_*, and minimize over both *x_s_* and these auxiliary variables.

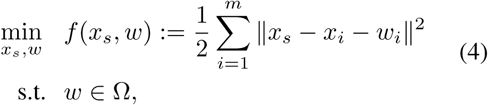

with Ω a special set described below. The original sets defined by (2) and (4) are both subset of ℝ^3^. The set describing (2) is a simple subset of ℝ^3^*^m^*:

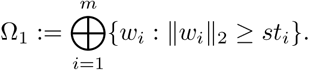

The set in (3) is also a subset of ℝ^3^*^m^*, with more complex structure:

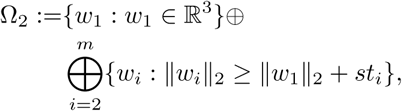

where ⊕ denote the direct sum. The original problem for calculating a projection onto a nonconvex set is difficult. By relaxing the formulation, the objective becomes more tractable, yielding a simple update rule using the structure of Ω. We also have a guarantee of optimality for the original feasibility problem based on the objective value of (4):

- Any *x_s_* satisfying (2) gives a global minimizer of (4) with objective value 0, by 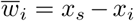.
- Any solution with zero objective value gives 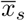 feasible with respect to (2) or (3).

Problem (4) may have a nonzero optimal value, in which case the ‘relaxed’ solution *x_s_* will not satisfy the original formulation. However, in practice we find a feasible point in each iteration. To solve (4), we minimize over *x_s_* and *w_i_*. Given {*w_i_*}, we have a closed for solution for *x_s_*:

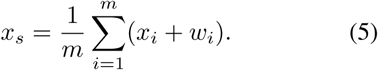

To find {*w_i_*} given *x_s_*, we have to solve

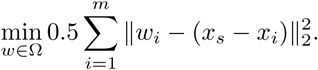

When Ω = Ω_1_, we have a closed form solution for the projection problem:

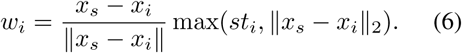

When Ω = Ω_2_, the projection is found by solving 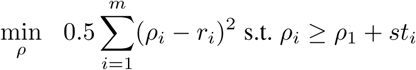, *i* ≥ 2, which requires a specialized subroutine. The approach is summarized in Algorithm 1.

#### Algorithm 1

Source Finding Algorithm

1. **Input**: {*x_i_*}, *s*
2. **Initialize**: *k* = 0, 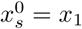, 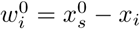
3. **while** not converged **do**
4. 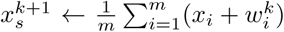
5. 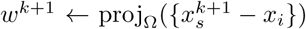
6. *k* ← *k* + 1
7. **Output**: 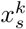

Algorithm 1 terminates when the function value or step-size is less than a specified tolerance, or if we hit an iteration cap of 200. It is equivalent to proximal gradient descent on the value function for (4):

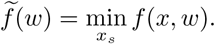

See [6] for an analysis of such algorithms, including rates of convergence.

**Robust modification**. The data collection process by a diagnostic catheter is inherently noisy, and some trial points give anomalous timing data. These anomalies then give incorrect information about the feasibility region (2). To make the method more robust, we detect and remove potential outliers in the course of solving each optimization problem (4). The outliers naturally give constraints that are very hard to satisfy. We introduce a vector τ to indicate which constraints are easy to fit, and which are difficult. The few constraints that are the least consistent with the remaining data are likely outliers. This idea can be traced back to least trimmed squares [3]; see also [1] for a survey of modern applications.

To implement the approach, we remove a small number h of arrival times from consideration in each iteration. We sort {*r*_1_ − *st*_1_*,…,r_m_ − st_m_*} from least to greatest, and for each index *i* in the smallest *h* residuals, we set *τ_i_* = 0, while all remaining *τ_i_* are set to 1. When we update *x_s_*, we modify (5) to 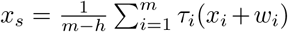. This strategy removes the influence of the potential outliers, and decreases the number of touches we need to find the source.

**Online implementation**. We start with several measurements obtained by a preliminary diagnostic catheter with 10 poles, see Figure 3. This sets up the first feasibility problem (2) or (3), which we solve by the reformulation (4) point to sample.

As we proceed, we consider the last 10 observations in forming each subsequent feasibility problem. If we accumulate a lot of data far away from the source, the feasibility problem becomes harder to solve; in particular the simple approximation of assuming a constant propagation speed s between the potential source *x_s_* and all observations is not reasonable. Algorithm 1 typically finds a solution (i.e. the next potential *x_s_* given current data) within 1-2 iterations for most problems.

## IV. Results

We run a simulation using real patient data. Timing and mesh data for three patients (two ventricular chambers and one atrium) are available from a diagnostic study, see Figures 2-5. The ‘ground truth’ of the arrhythmia source is inferred by the earliest arrival time observed during the entire data collection process. We start with 10 observations made by a diagnostic catheter, and use the online version of the algorithm to locate the source. Monte Carlo sampling is used to randomly initialize the initial 10 readings; we track the distance of estimates to source as a function of touches, and report the results across the simulations by using median, 75th, and 90th quantiles of the distance to source (mm) as a function of touches. A sample run of the algorithm is shown in Figure 3. The algorithm proposes the next point to sample after each touch. We then sample the closest point with data to each *x_s_* proposed by the algorithm; in practice the operator can simply attempt to move the catheter to a proposed point to obtain the next measurement. The procedure takes 12 touches to locate the true source for this run. For each touch, we need 1 or 2 iterations of Algorithm 1 to solve the feasibility problem. The objective values are numerically 0, which means each *x_s_* we find satisfies (2).

**Fig. 2.**
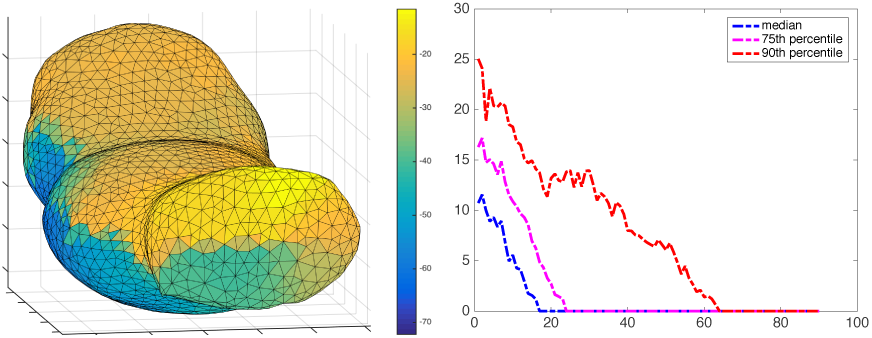
Left: simple ventricle dataset with color-coded arrival times. Right: distance to source as a function of touches using simple feasible region (2). In this simple dataset, the proposed approach quickly finds the source.

**Fig. 3.**
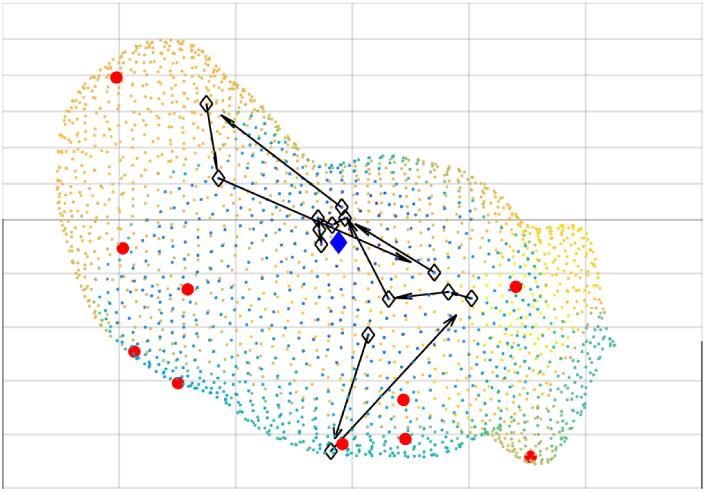
Path to source for sample run using dataset in Figure 2. Initial placement of diagnostic catheter is shown by red dots; path to source is shown using diamond markers.

**Fig. 4.**
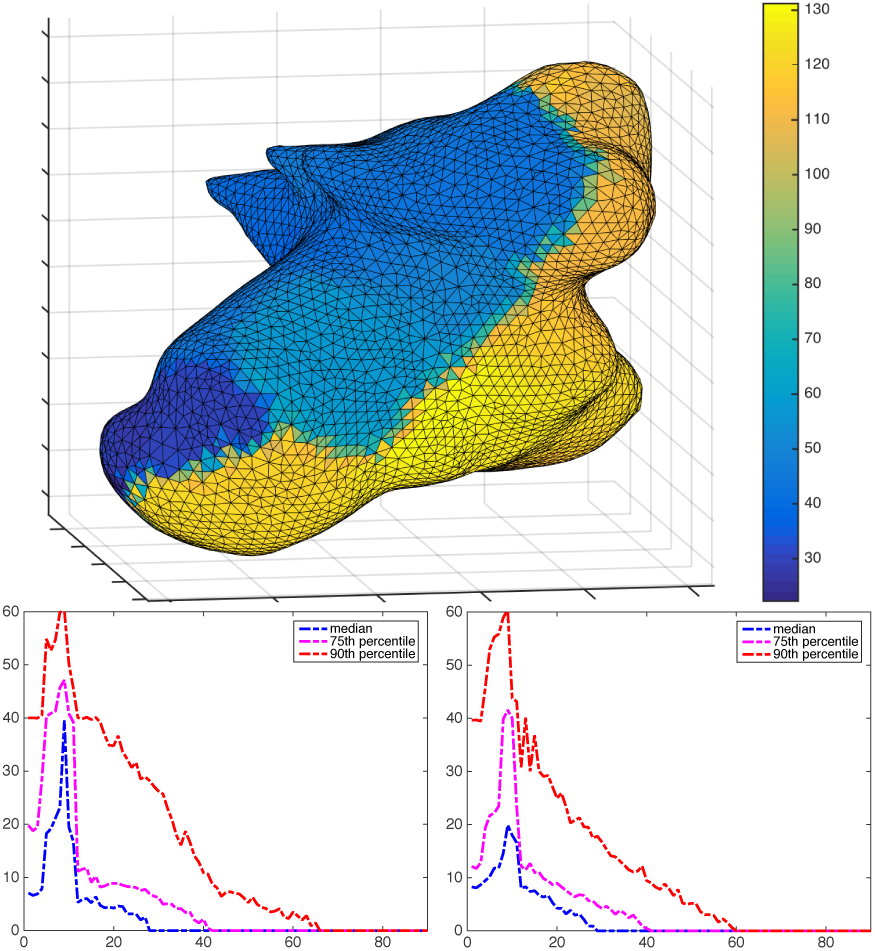
Top panel: ventricle dataset with color-coded arrival times. Bottom left: distance to source as a function of touches using simple feasible region (2). Bottom right: distance to source as a function of touches using coupled feasible region (3). Both approaches perform similarly on this example.

**Fig. 5.**
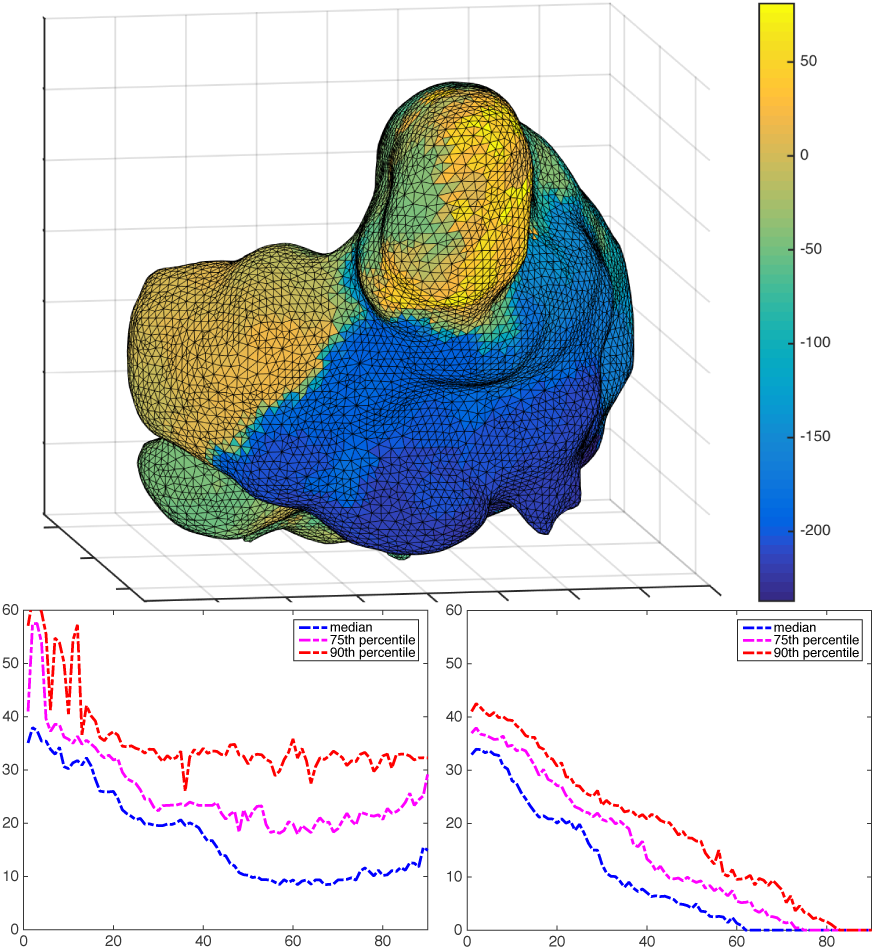
Top panel: atrial dataset with color-coded arrival times. Bottom left: distance to source as a function of touches using simple feasible region (2). Bottom right: distance to source as a function of touches using coupled feasible region (3). Accounting for coupling in the signal propagation gives a significant improvement on this challenging dataset.

Simulation results are plotted for each of the three datasets in Figures 2-5. For the first two datasets, using the simple feasible region (2) works as well as using the coupled encoding (3). For the more complex atrial dataset in Figure 5, adding more geometrical constraints pays off, and the results of (3) are significantly better. In all cases, Algorithm 1 easily solves the nonconvex feasibility problems required to tell the operator where to sample next.

## V. Conclusion

We formulated the arrhythmia localization problem as a sequence of nonconvex feasibility problems, and developed an efficient algorithm to solve these problems. The approach can accommodate different physical models of signal propagation in the heart; we compared two different models in this paper. The resulting approach opens a path to a new computationally-guided clinical approach to localize point-source arrhythmias, making online suggestions based on minimal information: we assume no prior knowledge about the heart’s anatomy. The next steps are to develop and test in the clinical setting with existing arrhythmia mapping systems.

## VI. Acknowledgements

We thank Boston Scientific, Inc for providing deidentified Rhythmia^®^ data sets to support this work. Dr. Aravkin was supported by the WRF Data Science Professorship.

## Footnotes

1 Boston Scientific, Marlborough, MA, USA.

